# Replicability of Structural Brain Alterations Associated with General Psychopathology: Evidence from a Population-Representative Birth Cohort

**DOI:** 10.1101/667220

**Authors:** Adrienne L. Romer, Annchen R. Knodt, Maria L. Sison, David Ireland, Renate Houts, Sandhya Ramrakha, Richie Poulton, Ross Keenan, Tracy R. Melzer, Terrie E. Moffitt, Avshalom Caspi, Ahmad R. Hariri

## Abstract

Transdiagnostic research has identified a general psychopathology factor – often called the ‘*p*’ factor – that accounts for shared variation across internalizing, externalizing, and thought disorders in diverse samples. It has been argued that the *p* factor may reflect dysfunctional thinking present in serious mental illness. In support of this, we previously used a theory-free, data-driven multimodal neuroimaging approach to find that higher *p* factor scores are associated with structural alterations within a cerebello-thalamo-cortical circuit (CTCC) and visual association cortex, both of which are important for monitoring and coordinating information processing in the service of executive control. Here we attempt to replicate these associations by conducting region-of-interest analyses of CTCC and visual association cortex using data from 875 members of the Dunedin Longitudinal Study, a five-decade study of a population-representative birth cohort now 45 years old. We further sought to replicate a more recent report that *p* factor scores can be predicted by patterns of distributed cerebellar morphology as estimated through independent component analysis. We successfully replicated associations between higher *p* factor scores and both reduced grey matter volume of the visual association cortex and fractional anisotropy of pontine white matter pathways within the CTCC. In contrast, we failed to replicate prior associations between cerebellar structure and *p* factor scores. Collectively, our findings encourage further focus on the CTCC and visual association cortex as core neural substrates and potential biomarkers of general psychopathology.

## Introduction

A rapidly emerging body of research has identified a general factor that captures shared variation among multiple forms of psychopathology across diverse samples (1). This general psychopathology or ‘*p*’ factor (2) accounts for the high rates of comorbidity among internalizing, externalizing, and thought disorders. Multiple explanations of the meaning of the *p* factor have been proposed, including that the *p* factor may index functional impairment, negative affect, emotion dysregulation, and poor intellectual function (for a review see (3)). One compelling argument regarding the nature of the *p* factor is that it captures the extent of disordered or dysfunctional thinking present not only in thought disorders, but also in extreme presentations of internalizing and externalizing disorders (3). Consistent with this argument, we recently used a theory-free, data-driven approach to find that among 1246 university students higher *p* factor scores were associated with structural alterations in a cerebello-thalamo-cortico circuit (CTCC) critical for monitoring and coordinating information processing in the service of executive control (4).

Specifically, we found that higher *p* factor scores were associated with reduced grey matter volume (GMV) in neocerebellar lobule VIIb. This neocerebellar region is a component of a specific CTCC, including the orbitofrontal, dorsolateral, and medial prefrontal cortices (5,6). We also found evidence for decreased microstructural integrity of pontine white matter pathways, as indexed by decreased fractional anisotropy (FA), which mediate communication of information from the prefrontal cortex to the neocerebellum within this CTCC (7–11). Investigators have theorized that this prefrontal CTCC plays a crucial role in comparing intention with the execution of thoughts, emotions, and actions by continuously updating internal models (12,13). Moreover, prefrontal CTCC dysfunction has been consistently reported in disorders principally characterized by poor executive control and disorganized thought such as schizophrenia (e.g., (14,15)), and individuals with cerebellar cognitive affective syndrome following damage to the neocerebellum experience executive control dysfunction symptoms referred to as “dysmetria of thought” (16–18).

A subsequent report based on analyses of data from 1401 community volunteers revealed that patterns of distributed cerebellar morphology also were associated with general psychopathology as estimated through independent component analysis (19). Namely, morphological features within a cerebellar component involved in cognitive functions (i.e., verbal working memory, retrieval, rehearsal, etc.), as well as reduced GMVs within neocerebellar lobule VI and crus I, were associated with higher general psychopathology. Further, these neocerebellar morphological features were the most important predictor of general psychopathology as compared to 52 other brain-wide anatomical features.

In addition to these structural alterations within neocerebellum and broader prefrontal CTCC, we found novel evidence for decreased GMV in the visual association cortex of individuals with higher *p* factor scores (4). Subsequently, we found that higher *p* factor scores were associated with patterns of inefficient intrinsic functional connectivity between visual association cortex and networks supporting executive control and self-referential processes, which are implicated across mental disorders (20). Collectively, these patterns are consistent with speculation that higher *p* factor scores ultimately represent the likelihood of experiencing disordered thought through a diminished capacity for basic monitoring and processing of information supported by the prefrontal CTCC and connectome-wide intrinsic functional connectivity. Such patterns of brain dysfunction may also contribute to negative affect, emotion dysregulation, and inefficient information processing, all of which also have been posited as potential explanations of the *p* factor (3).

It is important to seek to replicate these associations, especially because our original associations were discovered in a convenience sample of high-functioning 18 to 22 year-old university students through the Duke Neurogenetics Study (4). Here we attempt to replicate our original associations between prefrontal CTCC and visual association cortex structure and *p* factor scores using data from the Dunedin Longitudinal Study, a five-decade longitudinal study of a population-representative birth cohort now in midlife. Using data from the Dunedin Study, we further sought to replicate the independent components analysis of cerebellar morphology and general psychopathology as reported by Moberget et al. (19) in their study of young community volunteers.

## Materials and Methods

### Participants

Participants are members of the Dunedin Study, a longitudinal investigation of health and behavior in a representative birth cohort. Study members (n=1037; 91% of eligible births; 52% male) were all individuals born between April 1972 and March 1973 in Dunedin, New Zealand (NZ), who were eligible based on residence in the province and who participated in the first assessment at age 3 years (21). The cohort represented the full range of socioeconomic status (SES) in the general population of NZ’s South Island and as adults matched the NZ National Health and Nutrition Survey on key adult health indicators (e.g., body mass index, smoking, GP visits) and the NZ Census of citizens of the same age on educational attainment. Study members are primarily white (93%), matching South Island demographics (21). Assessments were carried out at birth and ages 3, 5, 7, 9, 11, 13, 15, 18, 21, 26, 32, 38, and most recently (completed April 2019) 45 years, when 94.1% (n=938) of the 997 participants still alive took part, and 875 (93%) of these age-45 participants also completed MRI scanning (see Supplementary Information, including Figure 1, for further details). Attrition analyses revealed that scanned Study members did not differ from other living Study members on *p* factor scores, childhood SES, or childhood IQ (see Supplementary Information for details). The relevant ethics committees approved each phase of the Study and informed consent was obtained from all Study members.

**Figure 1.**
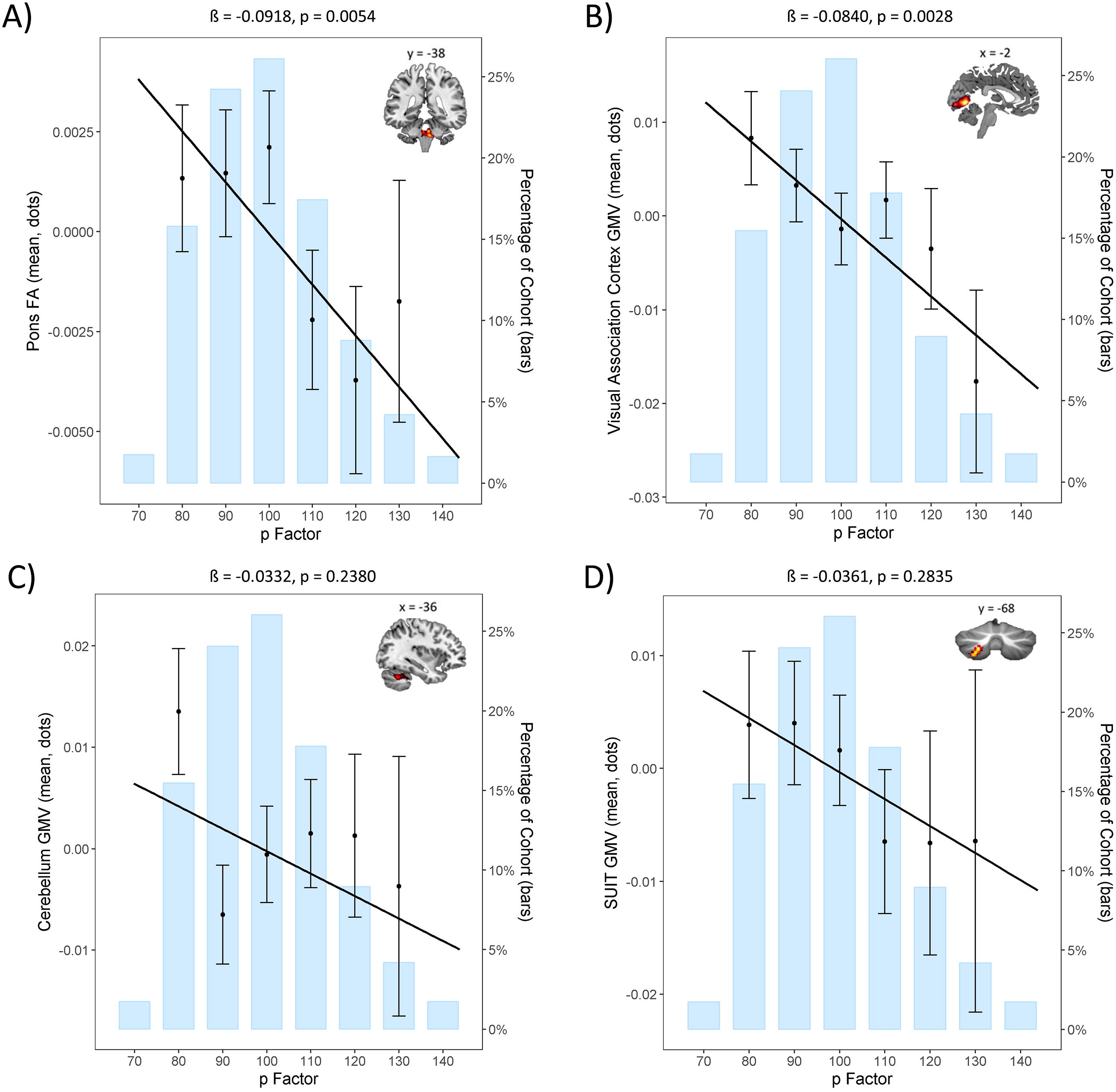
Replication analyses in the Dunedin Study of the original structural brain associations with *p* factor scores from Romer et al. (4). A) Replication of the negative association between pontine fractional anisotropy (FA) and *p* factor scores. B) Replication of the negative association between visual association cortex grey matter volume (GMV) and *p* factor scores. C) Non-significant replication of the negative association between cerebellar GMV and *p* factor scores. D) Non-significant replication of the negative association between SUIT-based neocerebellar lobule VIIb GMV and *p* factor scores. Per convention, *p* factor scores are normalized to a mean of 100 (SD = 15).

### Measuring the General Factor of Psychopathology, the p factor

The Dunedin Study longitudinally ascertains mental disorders every few years, interviewing members about past-year symptoms (see Supplementary Information, including Figure 2, for details). We studied Diagnostic and Statistical Manual of Mental Disorders (DSM)–defined symptoms of the following disorders that were repeatedly assessed in our longitudinal study: ADHD, alcohol dependence, cannabis dependence, dependence on hard drugs, tobacco dependence (assessed with the Fagerström Test for Nicotine Dependence (22)), conduct disorder, major depression, generalized anxiety disorder, fears and/or phobias, eating disorders, PTSD, obsessive compulsive disorder, mania, as well as positive and negative schizophrenia symptoms. Ordinal measures represented the number of possible DSM-defined symptoms associated with each disorder. Fears and/or phobias were assessed as the count of diagnoses for simple phobia, social phobia, agoraphobia, and panic disorder that a study member reported at each assessment. Symptoms were assessed without regard for hierarchical exclusionary rules to facilitate the examination of comorbidity. Each of the 14 disorders were assessed at least 3 times. The past-year prevalence rates of psychiatric disorders in the Dunedin cohort are similar to prevalence rates in nationwide surveys of the United States and New Zealand (23,24).

**Figure 2.**
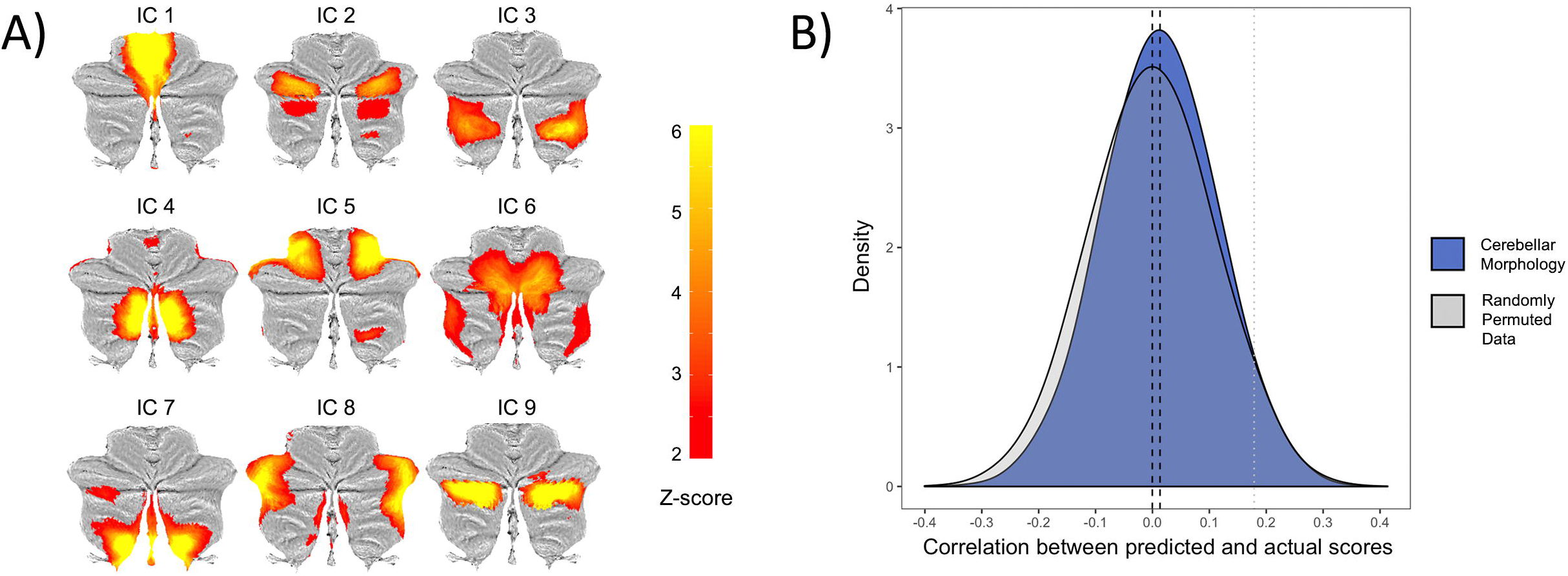
Replication analyses in the Dunedin Study of the original ICA-derived cerebellar morphology associations with *p* factor scores from Moberget et al. (19). A) The nine independent components resulting from data-driven decomposition of cerebellar grey matter maps projected onto flat-maps of the cerebellar cortex (36). B) Distributions of correlations between predicted and actual *p* factor scores across 10,000 iterations of the 10-fold cross-validated model using the average of the 9 independent components from A compared to the empirical null-distribution. The black dotted lines represent the mean for each distribution and the grey dotted line represents the one-tailed .05 threshold. The nine ICA-derived components predicted *p* factor scores beyond chance on average, but the difference from the empirical null distribution was p = 0.53, suggesting non-significant replication of Moberget et al. (19).

The method used to compute a general factor of psychopathology in the Dunedin cohort up to age 38 has been described previously (2); here we extend these models to include the age-45 data (see Supplementary Information for details). Briefly, we used confirmatory factor analysis to compute a bi-factor model specifying a general psychopathology factor (labeled *p*) (Supplementary Figure 3). This model included our 14 observed variables: ADHD, alcohol dependence, cannabis dependence, dependence on hard drugs, tobacco dependence, conduct disorder, major depression, generalized anxiety disorder, fears/phobias, eating disorders, PTSD, obsessive-compulsive disorder, mania, as well as positive and negative schizophrenia symptoms. The model also included method/state factors designed to pull out age-and assessment-related variance (e.g., interviewer effects, mood effects, and age-specific vulnerabilities) that was uncorrelated with trait propensity toward psychopathology.

All analyses were performed in Mplus version 7.12 using the weighted least squares means and variance adjusted (WLSMV) algorithm. After respecification for a Heywood case, the bi-factor model fit the data well (Supplementary Table 1 and Figure 3): χ^2^(2457, N = 1000) = 3695.364, CFI = .949, TLI = .945, RMSEA = .022, 90% confidence internal (CI) = [.021, .024]. Loadings on the *p* factor were high (all p’s<.001) and averaged .612. For expository purposes, we scaled Study members’ *p* factor scores to M=100, SD=15. The *p* factor allows us to test for structural brain alterations in relation to general psychopathology. Study members with higher *p* factor scores experienced a greater variety of mental disorders from adolescence to midlife (r=.77; Supplementary Figure 4).

### MRI Data Acquisition

Each study member was scanned using a Siemens Skyra 3T scanner equipped with a 64-channel head/neck coil at the Pacific Radiology imaging center in Dunedin, New Zealand. Diffusion-weighted images providing full brain coverage were acquired with 2.5 mm isotropic resolution and 64 diffusion weighted directions (4700 ms repetition time, 110.0 ms echo time, b value 3,000 s/mm2, 240 mm field of view, 96×96 acquisition matrix, slice thickness = 2.5 mm). Non-weighted (b = 0) images were acquired in both the encoding (AP) and reverse encoding (PA) directions to allow for EPI distortion correction. High resolution structural images were obtained using a T1-weighted MP-RAGE sequence with the following parameters: TR = 2400 ms; TE = 1.98 ms; 208 sagittal slices; flip angle, 9°; FOV, 224 mm; matrix = 256×256; slice thickness = 0.9 mm with no gap (voxel size 0.9×0.875×0.875 mm); and total scan time = 6 min and 52 s. All neuroimaging data were visually inspected for quality. Data were excluded for Study members who were unable to be scanned with the 64-channel head coil, had an incidental finding, or whose scans were of poor quality due to motion (as revealed by visual inspection for T1-weighted images or >3 mm frame-to-frame movements for diffusion images), resulting in a total of 854 study members eligible for diffusion analyses and 860 study members eligible for GMV analyses.

### Fractional Anisotropy

Following the methods of Romer et al. (4), diffusion tensor imaging analyses were completed using SPM8 implemented in Matlab R2016a. All diffusion weighted scans were motion corrected and co-registered to the mean image to correct for head movement. The tensor model was used to calculate FA values for each voxel and non-brain tissue was removed. Each image was normalized to MNI space and smoothed using a 4 mm FWHM Gaussian kernel. We note that the tensor model for derivation of FA values is not optimized for our current diffusion-weighted image data (25), which was acquired with b = 3000 s/mm2 to facilitate future probabilistic tractography. We are unaware of any suitable alternatives for the derivation of FA values at higher b values. Moreover, these differences in acquisition parameters are of less concern because visual inspection of the preprocessed images revealed adequate registration and we did successfully replicate the association between higher *p* factor scores and lower pontine FA (see below).

### Grey Matter Volume

Again, following the methods of Romer et al. (4), regional GMVs were determined using the unified segmentation (26) and DARTEL normalization (27) modules in SPM12 (http://www.fil.ion.ucl.ac.uk/spm). Using this approach, individual T1-weighted images were segmented into grey, white, and CSF images, and then non-linearly registered to the existing IXI template of 550 healthy subjects averaged in standard Montreal Neurological Institute (MNI) space, available with VBM8 (http://dbm.neuro.uni-jena.de/vbm/). Subsequently, grey matter images were modulated for nonlinear effects of the high-dimensional normalization to preserve the total amount of signal from each region and smoothed with an 8 mm FWHM Gaussian kernel. The voxel size of processed images was 1.5×1.5×1.5 mm. A grey matter mask for subsequent analyses was created by thresholding the final stage (6th) IXI template at 0.1.

### Cerebellar Grey Matter Volume

In addition to the above whole-brain voxel based GMV analyses, the Spatially Unbiased Infratentorial Toolbox (SUIT) was used for high-resolution cerebellar-specific voxel-based morphometry analyses per the methods of Romer et al. (4). For each Study member, the Isolate function of the toolbox was used to create a mask of the cerebellum and generate grey and white matter segmentation maps. The masked segmentation maps were then normalized to the SUIT template with non-linear DARTEL normalization. The resulting cerebellar grey matter image was resliced into the SUIT atlas space and smoothed with a 4 mm FWHM isotropic Gaussian kernel, a small kernel to preserve precision in the definition of cerebellar structures, in line with previous publications (28).

### Independent Component Analysis of Cerebellar Morphology

Lastly, we conducted an independent component analysis (ICA) of SUIT-based cerebellar morphology using the method of Moberget et al. (19). Briefly, we masked the SUIT-derived cerebellar grey matter maps using the SUIT toolbox’s grey matter probability map thresholded at 0.1 and subjected them to ICA using FSL MELODIC (29). In our sample, a model order of 9 corresponded to the highest number of clearly bilateral components, and this model was used for further analyses.

### Statistical Analyses

Exact masks were created from the three primary associations with *p* factor scores originally reported in Romer et al. (4): a 272 voxel cluster in the pons, a 2353 voxel cluster in the visual association cortex, and a 706 voxel cluster in the cerebellum. A fourth mask was created for the 156 voxel cluster in neocerebellar lobule VIIb identified through the SUIT analysis. Moving to the Dunedin Study data, mean values for each of these four masks were extracted for each Study member from the FA (pons), GMV (visual association cortex and neocerebellum), and SUIT maps, respectively. These mean extracted values were then used as the dependent variable in linear models with *p* factor scores as the predictor and sex and total intracranial volume or average total FA, respectively, as covariates to explicitly test for replication of the original findings of Romer et al. (4).

Per the strategy of Moberget et al. (19), we also tested whether weights on our nine ICA-derived cerebellar components could predict *p* factor scores using shrinkage linear regression with 10,000 iterations of 10-fold cross-validation on randomly partitioned data. As in Moberget et al. (19), we controlled for sex and total intracranial volume. Performance was evaluated by comparing the distribution of Pearson correlations between predicted and observed *p* factor scores to a null distribution of correlations obtained by randomly permuting the *p* factor scores.

## Results

### White Matter Microstructural Integrity

A significant negative correlation (standardized β = −0.092; p = 0.005) indicated an association between lower pontine FA and higher *p* factor scores (Figure 1A), replicating the finding of Romer et al. (4).

### Grey Matter Volume

A significant negative correlation (standardized β = −0.084; p = 0.003) indicated an association between decreased visual association cortex GMV and higher *p* factor scores (Figure 1B), replicating the finding of Romer et al. (4). An observed negative correlation between cerebellar GMV and *p* factor scores was not statistically significant (standardized β = −0.033; p = 0.238; Figure 1C). This was also true for the SUIT-based neocerebellar lobule VIIb cluster (standardized β = −0.036; p = 0.284; Figure 1D).

### ICA-Derived Cerebellar Morphology

The nine independent components of cerebellar morphology collectively accounted for 41.47% of the total variance in the modulated grey matter maps; each component explained between 4.08% and 4.97% of the total variance (and between 9.83% and 11.99% of the explained variance). The nine ICA-derived components predicted *p* factor scores beyond chance on average, but the difference from the empirical null distribution was not significant (mean correlations between predicted and observed values: r = 0.13, p = 0.53; mean r > 54.89% of the empirical null distribution; Figure 2).

## Discussion

We successfully replicated two prior associations between variation in brain structure and general psychopathology, as indexed by the *p* factor, using data from a population-representative birth cohort now in midlife. Namely, we replicated associations between *p* factor scores and both pontine FA and visual association cortex GMV as originally reported by Romer et al. (4). In contrast, we failed to replicate three prior associations between cerebellar structure and *p* factor scores. First, although nominally consistent with the original report of Romer et al. (4), neither of two tested associations between GMV in a broad cerebellar cluster nor a smaller cluster in neocerebellar lobule VIIb were statistically significant. Second, an ICA-based measure of global cerebellar morphology did not significantly predict *p* factor scores above chance as was reported originally by Moberget et al. (19).

The replication of a negative association between pontine FA and *p* factor scores further implicates the CTCC in general psychopathology. Thus, dysfunction in fundamental aspects of monitoring and coordinating executive functions (i.e., “forward control”) through dynamic information processing between the neocerebellum and prefrontal cortex appears to be a core transdiagnostic feature of general psychopathology. The second replication of a negative association between *p* factor scores and GMV in visual association cortex is consistent with the importance of executive dysfunction in general psychopathology. In particular, structural alterations in visual association cortex may manifest as more effortful or less efficient integration of bottom-up sensory information with attentional demands and executive control processes in individuals who meet criteria for different forms of mental disorders (20).

The non-significant associations between *p* factor scores and multiple indices of cerebellar GMV and morphology do not necessarily undermine the importance of a prefrontal CTCC in general psychopathology. Rather, these failures may indicate that shared variation among different forms of psychopathology, as captured by the *p* factor, is more a reflection of how information is communicated within the CTCC, particularly through pontine white matter pathways connecting the prefrontal cortex and cerebellum, and less a reflection of how information may be locally computed within the neocerebellum. This would be consistent with the emerging understanding that brain function may be best characterized by distributed patterns of network communication rather than discrete regional activity (30,31). However, there also are pragmatic factors that may have limited our ability to replicate prior associations between *p* factor scores and cerebellar structure.

First, the failure to replicate cerebellar associations with *p* factor scores may reflect different contributions of brain structure to risk across development. The discovery samples were comprised of young adults (4) or children, adolescents, and young adults (19). In contrast, our sample is comprised of individuals in midlife. Thus, the contribution of cerebellar GMV and morphology to the *p* factor may be greater earlier than later in life. This difference may reflect the still-active structural development of the cerebellum, which parallels that of the prefrontal cortex, in both discovery samples (32). Developmental differences are hinted at by the observation of only nine independent components of cerebellar morphology in our sample but ten such components in the sample studied by Moberget et al. (19). Longitudinal assessment of brain structure and *p* factor scores within the same individuals is necessary to evaluate a hypothesis of developmental differences (33). Second, the nature of the sampling strategy across the three samples also may influence replication. Unlike the population-representative birth cohort in our current study, both discovery samples represented narrow groups of select individuals (e.g., high-functioning university students or community volunteers). Additional replication efforts across diverse samples are necessary to probe the implications of such possible differences for the study of the brain basis of general psychopathology. Lastly, we may simply have been underpowered to identify significant associations of small effect. Our current sample is smaller than either the discovery sample of Romer et al. (4) (N=1246) or Moberget et al. (19) (N=1401). Generally, successful replication is more likely if the test samples are larger and thus better powered to detect often smaller effects than reported in a discovery sample (34,35). The effect sizes in Romer et al. (4) ranged from r = 0.09 -.13 and those in Moberget et al. (19) ranged from r = .13 -.2.

These limitations notwithstanding, the two replicable associations of the theory-free, data-driven findings of Romer et al. (4) reported herein point to specific features of brain structure that may be a core feature of shared variation among common forms of mental illness. Alterations in the microstructural integrity of pontine white matter pathways may reflect dysfunction of executive control processes supported through dynamic communication within the CTCC. Likewise, alterations in GMV of visual association cortex may reflect impairments in the integration of bottom-up sensory information with top-down executive control and attentional processes. Notably, both of these neuroanatomical features are consistent with a model of the *p* factor as indexing increasingly disordered thought, which characterizes the most debilitating forms of mental disorders. The extent to which these neuroanatomical features drive the emergence of general psychopathology or emerge as a consequence of general psychopathology are as yet unknown and require longitudinal neuroimaging assessments to explicate.

## Acknowledgments

The Dunedin Multidisciplinary Health and Development Study is supported by the NZ HRC and NZ MBIE. This research was supported by National Institute on Aging grant R01AG032282, R01AG049789, and UK Medical Research Council grant MR/P005918/1. Additional support was provided by the Jacobs Foundation. ALR received support from the National Science Foundation Graduate Research Fellowship under Grant No. DGE 1106401. Thank you to members of the Advisory Board for the Dunedin Neuroimaging Study. We thank the Dunedin Study members, unit research staff, Pacific Radiology staff, and Study founder Phil Silva.

## References

1. Lahey BB, Krueger RF, Rathouz PJ, Waldman ID, Zald DH. A hierarchical causal taxonomy of psychopathology across the life span. Psychol Bull. 2017 Feb;143(2):142–86.

2. Caspi A, Houts RM, Belsky DW, Goldman-Mellor SJ, Harrington H, Israel S, et al. The p factor: One general psychopathology factor in the structure of psychiatric disorders? Clin Psychol Sci J Assoc Psychol Sci. 2014 Mar;2(2):119–37.

3. Caspi A, Moffitt TE. All for one and one for all: mental disorders in one dimension. Am J Psychiatry. 2018 Apr 6;175(9):831–44.

4. Romer AL, Knodt AR, Houts R, Brigidi BD, Moffitt TE, Caspi A, et al. Structural alterations within cerebellar circuitry are associated with general liability for common mental disorders. Mol Psychiatry. 2018 Apr;23(4):1084–90.

5. Buckner RL, Krienen FM, Castellanos A, Diaz JC, Yeo BTT. The organization of the human cerebellum estimated by intrinsic functional connectivity. J Neurophysiol. 2011 Nov;106(5):2322–45.

6. Thomas Yeo BT, Krienen FM, Sepulcre J, Sabuncu MR, Lashkari D, Hollinshead M, et al. The organization of the human cerebral cortex estimated by intrinsic functional connectivity. J Neurophysiol. 2011 Sep;106(3):1125–65.

7. Middleton FA, Strick PL. Cerebellar projections to the prefrontal cortex of the primate. J Neurosci. 2001 Jan 15;21(2):700–12.

8. Buckner RL. The cerebellum and cognitive function: 25 years of insight from anatomy and neuroimaging. Neuron. 2013 Oct 30;80(3):807–15.

9. D’Angelo E, Casali S. Seeking a unified framework for cerebellar function and dysfunction: from circuit operations to cognition. Front Neural Circuits [Internet]. 2013 [cited 2019 Aug 15];6. Available from: https://www.frontiersin.org/articles/10.3389/fncir.2012.00116/full#F3

10. Keren-Happuch E, Chen S-HA, Ho M-HR, Desmond JE. A meta-analysis of cerebellar contributions to higher cognition from PET and fMRI studies. Hum Brain Mapp. 2014 Feb;35(2):593–615.

11. Marek S, Siegel JS, Gordon EM, Raut RV, Gratton C, Newbold DJ, et al. Spatial and temporal organization of the individual human cerebellum. Neuron. 2018 Nov 21;100(4):977–993.e7.

12. Ito M. Movement and thought: identical control mechanisms by the cerebellum. Trends Neurosci. 1993 Nov 1;16(11):448–50.

13. Ito M. Control of mental activities by internal models in the cerebellum. Nat Rev Neurosci. 2008 Apr;9(4):304–13.

14. Bernard JA, Mittal VA. Dysfunctional activation of the cerebellum in schizophrenia: A functional neuroimaging meta-analysis. Clin Psychol Sci J Assoc Psychol Sci. 2015 Jul 1;3(4):545–66.

15. Moberget T, Doan NT, Alnæs D, Kaufmann T, Córdova-Palomera A, Lagerberg TV, et al. Cerebellar volume and cerebellocerebral structural covariance in schizophrenia: a multisite mega-analysis of 983 patients and 1349 healthy controls. Mol Psychiatry. 2018 Jun;23(6):1512–20.

16. Andreasen NC, Paradiso S, O’Leary DS. “Cognitive Dysmetria” as an integrative theory of schizophrenia: A dysfunction in cortical-subcortical-cerebellar circuitry? Schizophr Bull. 1998 Jan 1;24(2):203–18.

17. Schmahmann JD. Disorders of the cerebellum: Ataxia, dysmetria of thought, and the cerebellar cognitive affective syndrome. J Neuropsychiatry Clin Neurosci. 2004 Aug 1;16(3):367–78.

18. Schmahmann JD, Weilburg JB, Sherman JC. The neuropsychiatry of the cerebellum — insights from the clinic. The Cerebellum. 2007 Sep 1;6(3):254–67.

19. Moberget T, Alnæs D, Kaufmann T, Doan NT, Córdova-Palomera A, Norbom LB, et al. Cerebellar gray matter volume is associated with cognitive function and psychopathology in adolescence. Biol Psychiatry. 2019 Jul 1;86(1):65–75.

20. Elliott ML, Romer A, Knodt AR, Hariri AR. A Connectome-wide functional signature of transdiagnostic risk for mental illness. Biol Psychiatry. 2018 Sep 15;84(6):452–9.

21. Poulton R, Moffitt TE, Silva PA. The Dunedin Multidisciplinary Health and Development Study: Overview of the first 40 years, with an eye to the future. Soc Psychiatry Psychiatr Epidemiol. 2015 May 1;50(5):679–93.

22. Heatherton TF, Kozlowski LT, Frecker RC, Fagerström KO. The Fagerström Test for Nicotine Dependence: a revision of the Fagerström Tolerance Questionnaire. Br J Addict. 1991 Sep;86(9):1119–27.

23. Schaefer JD, Caspi A, Belsky DW, Harrington H, Houts R, Horwood LJ, et al. Enduring mental health: Prevalence and prediction. J Abnorm Psychol. 2017 Feb;126(2):212–24.

24. Moffitt TE, Caspi A, Taylor A, Kokaua J, Milne BJ, Polanczyk G, et al. How common are common mental disorders? Evidence that lifetime prevalence rates are doubled by prospective versus retrospective ascertainment. Psychol Med. 2010 Jun;40(6):899–909.

25. Steven AJ, Zhuo J, Melhem ER. Diffusion Kurtosis Imaging: An emerging technique for evaluating the microstructural environment of the brain. Am J Roentgenol. 2013 Dec 26;202(1):W26–33.

26. Ashburner J, Friston KJ. Unified segmentation. NeuroImage. 2005 Jul 1;26(3):839–51.

27. Ashburner J. A fast diffeomorphic image registration algorithm. NeuroImage. 2007 Oct 15;38(1):95–113.

28. D’Agata F, Caroppo P, Boghi A, Coriasco M, Caglio M, Baudino B, et al. Linking coordinative and executive dysfunctions to atrophy in spinocerebellar ataxia 2 patients. Brain Struct Funct. 2011 Sep 1;216(3):275–88.

29. Beckmann CF, Smith SM. Probabilistic independent component analysis for functional magnetic resonance imaging. IEEE Trans Med Imaging. 2004 Feb;23(2):137–52.

30. Mišić B, Sporns O. From regions to connections and networks: new bridges between brain and behavior. Curr Opin Neurobiol. 2016 Oct 1;40:1–7.

31. Bassett DS, Sporns O. Network neuroscience. Nat Neurosci. 2017 Feb 23;20(3):353–64.

32. Diamond A. Close interrelation of motor development and cognitive development and of the cerebellum and prefrontal cortex. Child Dev. 2000 Jan;71(1):44–56.

33. Schaie K. Age changes and age differences. The Gerontologist. 1967;7:128–32.

34. Anderson SF, Maxwell SE. Addressing the “Replication Crisis”: Using original studies to design replication studies with appropriate statistical power. Multivar Behav Res. 2017 May 4;52(3):305–24.

35. Turner BO, Paul EJ, Miller MB, Barbey AK. Small sample sizes reduce the replicability of task-based fMRI studies. Commun Biol. 2018 Jun 7;1(1):1–10.

36. Diedrichsen J, Zotow E. Surface-Based Display of Volume-Averaged Cerebellar Imaging Data. PLOS ONE. 2015 Jul 31;10(7):e0133402.

