## Supplemental Information for "Replicability of Structural Brain Alterations Associated with General Psychopathology: Evidence from a Population-Representative Birth Cohort"

**Supplementary Information**

*Supplemental Phase 45 Attrition Analysis*

We conducted an attrition analysis (**Supplementary Figure 1**) using *p* factor scores, childhood intelligence quotient (IQ), and socioeconomic status (SES) to determine whether participants in the Phase 45 data collection were representative of the original cohort. We found no significant differences in *p* factor scores between the full cohort, those still alive, those seen at Phase 45 or those scanned at Phase 45. Those who were deceased by the Phase 45 data collection had significantly higher *p* factor scores than those who were still alive (t = -2.86, p = 0.004). No significant differences in childhood IQ were found between the full cohort, those still alive, those seen at Phase 45 or those scanned at Phase 45. Those who were deceased by the Phase 45 data collection had significantly lower childhood IQ’s than those who were still alive (t = 2.09, p = 0.04). No significant differences were found between the full cohort, those deceased, those alive, those seen at Phase 45 or those scanned at Phase 45 on childhood SES.

*Assessing Psychopathology*

Mental disorders are disturbances in thought, behavior, and emotion that interfere with or limit social, family, educational, or work activities. In the Dunedin Study, these were identified according to the criteria of the Diagnostic and Statistical Manual of Mental Disorders (DSM).

Psychiatric interviews were carried out by health professionals, not lay interviewers. At ages 18, 21, 26, 32, 38, and 45, interviews were carried out with the Diagnostic Interview Schedule (1) (2). The following disorders were assessed (**Supplementary Figure 2**): Externalizing (ADHD, Conduct Disorder, Alcohol Dependence, Tobacco Dependence, Cannabis Dependence, Other Drug Dependence), Internalizing (Generalized Anxiety Disorder, Depression, Fears [including Social Phobia, Simple Phobia, Agoraphobia, Panic Disorder], Eating Disorders [including Bulimia and Anorexia], PTSD), and Thought disorders (Obsessive-Compulsive Disorder, Mania, Schizophrenia). Diagnoses ignored hierarchical exclusionary rules to facilitate study of comorbidity. Diagnoses were based on symptom algorithms and impairment ratings, but also incorporated additional information including standardized teacher/parent/informant reports as developmentally appropriate, psychiatrists’ review of interviewers’ detailed case notes, pharmacists’ medication review, and staff ratings of symptoms observed (3).

At ages 18 and 21, diagnoses were made according to DSM-III-R (4); at ages 26, 32, and 38, according to DSM-IV (5); at age 45 according to the now-current DSM-V (6) (with the exception of substance-dependence disorders which were diagnosed according to DSM-IV, given that DSM-V dropped the distinction between abuse and dependence).

*Modelling the Structure of Psychopathology*

We have previously described the structure of psychopathology up to age 38 years (7); here we extend these models to include the age 45 data.

To evaluate the structure of psychopathology we used data from 6 adult assessments, carried out at ages 18, 21, 26, 32, 38, and 45 years. We studied DSM-defined symptoms of the following disorders that were repeatedly assessed in our longitudinal study: ADHD, Conduct Disorder, Alcohol Dependence, Cannabis Dependence, Dependence on Hard Drugs, Tobacco Dependence (assessed with the Fagerström Test for Nicotine Dependence (8)), Depression, Generalized Anxiety Disorder, Fears/Phobias (Social Phobia, Simple Phobia, Agoraphobia, Panic Disorder), PTSD, Eating Disorders (Anorexia, Bulimia), Obsessive-Compulsive Disorder, Mania, as well as positive and negative Schizophrenia symptoms. Ordinal measures represented the number of the observed DSM-defined symptoms associated with each disorder. Fears/phobias were assessed as the count of diagnoses for simple phobia, social phobia, agoraphobia, and panic disorder that a study member reported at each assessment. Of the 14 disorders, 6 were not assessed at every occasion, but each disorder was measured at least three times (**Supplementary** **Figure 2**). Of the original 1,037 study members, we included 1,000 study members who had symptom count assessments for at least one age (845 study members had present symptom counts for all six assessments, 90 for five, 30 for four, 13 for three, and 14 for two). The 37 excluded study members comprised those who died (N=13) or left the study (N=21) before age 18 or who had such severe developmental disabilities (N=3) that they could not be interviewed with the Diagnostic Interview Schedule.

Using Confirmatory Factor Analysis (CFA), we tested a hierarchical or bifactor model that is frequently used to examine the structure of psychopathology (9). In CFA, latent continuous factors are hypothesized to account for the pattern of covariance among observed variables. Our CFA was run as multitrait-multimethod model. Observed variables represented each of the disorders with a symptom scale at each assessment age (e.g., alcohol dependence was measured with a symptom scale at ages 18, 21, 26, 32, 38, and 45). The model also included method/state factors designed to pull out age- and assessment-related variance (e.g., interviewer effects, mood effects, and age-specific vulnerabilities) that was uncorrelated with trait propensity toward psychopathology. Because symptom-level data are ordinal and have highly skewed distributions, we used polychoric correlations. Polychoric correlations provide estimates of the Pearson correlation by mapping thresholds to underlying normally distributed continuous latent variables that are assumed to give rise to the observed ordinal variables. Analyses were performed in Mplus version 8.3 (10) using the weighted least squares means and variance adjusted (WLSMV) algorithm. The WLSMV estimator is appropriate for categorical and nonmultivariate normal data and provides consistent estimates when data are missing at random with respect to covariates (11). We assessed how well the model fit the data using the chi-square value, the comparative fit index (CFI), the Tucker-Lewis index (TLI), and the root-mean-square error of approximation (RMSEA). CFI values greater than .95 and TLI values greater than 0.95 indicate good fit; RMSEA scores less than .05 are considered good (12).

The hierarchical or bifactor model (**Supplementary** **Figure 3**) tests the hypothesis that the symptom measures reflect both General Psychopathology and three narrower styles of psychopathology. General Psychopathology (labeled *p* in **Supplementary Figure 3**) is represented by a factor that directly influences all of the diagnostic symptom factors. In addition, styles of psychopathology are represented by three factors, each of which influences a smaller subset of the symptom items. For example, alcohol symptoms load jointly on the General Psychopathology factor and on the Externalizing style factor. The specific factors represent the constructs of Externalizing, Internalizing, and Thought Disorder over and above General Psychopathology. The model had a Heywood case, an estimated variance that was negative for one of the lower-order disorder/symptom factors (specifically, mania), suggesting this was not a valid model. Inspection of the results revealed the source of the convergence problem. Specifically, the Thought Disorder factor was subsumed in *p*; that is, in the hierarchical model, symptoms of OCD, mania, and schizophrenia loaded very highly on *p*, but unlike symptoms of Externalizing and Internalizing, they could not form a separate Thought Disorder factor independently of *p*. We respecified the model accordingly, depicted as the second in **Supplementary Figure 3**. This model fit the data well: χ^2^(2457, N=1,000) = 3695.364, CFI = .949, TLI = .945, RMSEA = .022, 90% CI [.021, .024]. As shown in **Supplementary Table 1**, loadings on the General factor (*p*) were all positive, generally high (all ps < .001), and averaged .612; the highest standardized loadings were for mania (.976), schizophrenia (.865), PTSD (.860), and OCD (.772). **Supplementary Figure 4** shows that the *p* factor captures how cohort members differ from each other in the variety and persistence of many different kinds of disorders over the adult life course. Cohort members with higher *p* factor scores experienced a greater variety of psychiatric disorders from adolescence to midlife (r=.77).

**Figure Captions**

**Supplementary Figure 1.** Attrition analysis using *p* factor scores and childhood IQ and SES shows that Study members who completed the neuroimaging protocol were representative of the original cohort.

No significant differences in *p* factor scores (top panel) were found between the full cohort, those still alive, those seen at Phase 45 or those scanned at Phase 45. Those who were deceased by the Phase 45 data collection had significantly higher *p* factor scores than those who were still alive (t = -2.86, p = 0.004). No significant differences in childhood IQ (middle panel) were found between the full cohort, those still alive, those seen at Phase 45 or those scanned at Phase 45. Those who were deceased by the Phase 45 data collection had significantly lower childhood IQ’s than those who were still alive (t = 2.09, p = 0.04). No significant differences were found between the full cohort, those deceased, those alive, those seen at Phase 45 or those scanned at Phase 45 on childhood SES (bottom panel).

**Supplementary Figure 2.** Structure of mental disorder data collected in the Dunedin Study.

The chart shows the age at which each disorder was assessed, from ages 18-45 years. Although each disorder was not assessed at every age, each disorder was assessed on at least three occasions.

**Supplementary Figure 3.** The structure of psychopathology.

Using Confirmatory Factor Analysis (CFA), we tested a hierarchical or bifactor model. Using this model, we tested the hypothesis that the symptom measures reflect both General Psychopathology and three narrower styles of psychopathology. General Psychopathology (labeled *p*) is represented by a factor that directly influences all of the diagnostic symptom factors. In addition, styles of psychopathology are represented by three factors, each of which influences a smaller subset of the symptom items. For example, alcohol symptoms load jointly on the General Psychopathology factor and on the Externalizing style factor. The specific factors represent symptoms of Externalizing, Internalizing, and Thought Disorder that are independent of General Psychopathology. The model had a Heywood case, an estimated variance that was negative for one of the lower-order disorder/symptom factors (specifically, mania), suggesting this was not a valid model. Inspection of the results revealed the source of the convergence problem. Specifically, the Thought Disorder factor was subsumed in *p*; that is, in the hierarchical model, symptoms of OCD, mania, and schizophrenia loaded very highly on *p*, but unlike symptoms of Externalizing and Internalizing, they could not form a separate Thought Disorder factor independently of *p*. We respecified the model accordingly, as shown in the second diagram.

**Supplementary Figure 4.** Plot of the positive correlation between the variety and persistence of mental disorders and *p* factor scores in the Dunedin Study.

The *p* factor captures how Study members differ from each other in the variety and persistence of many different kinds of disorders over the life course. Study members with higher *p* factor scores experienced a greater variety of mental disorders from adolescence to midlife (r=.77).

**Supplementary Figure 1.**

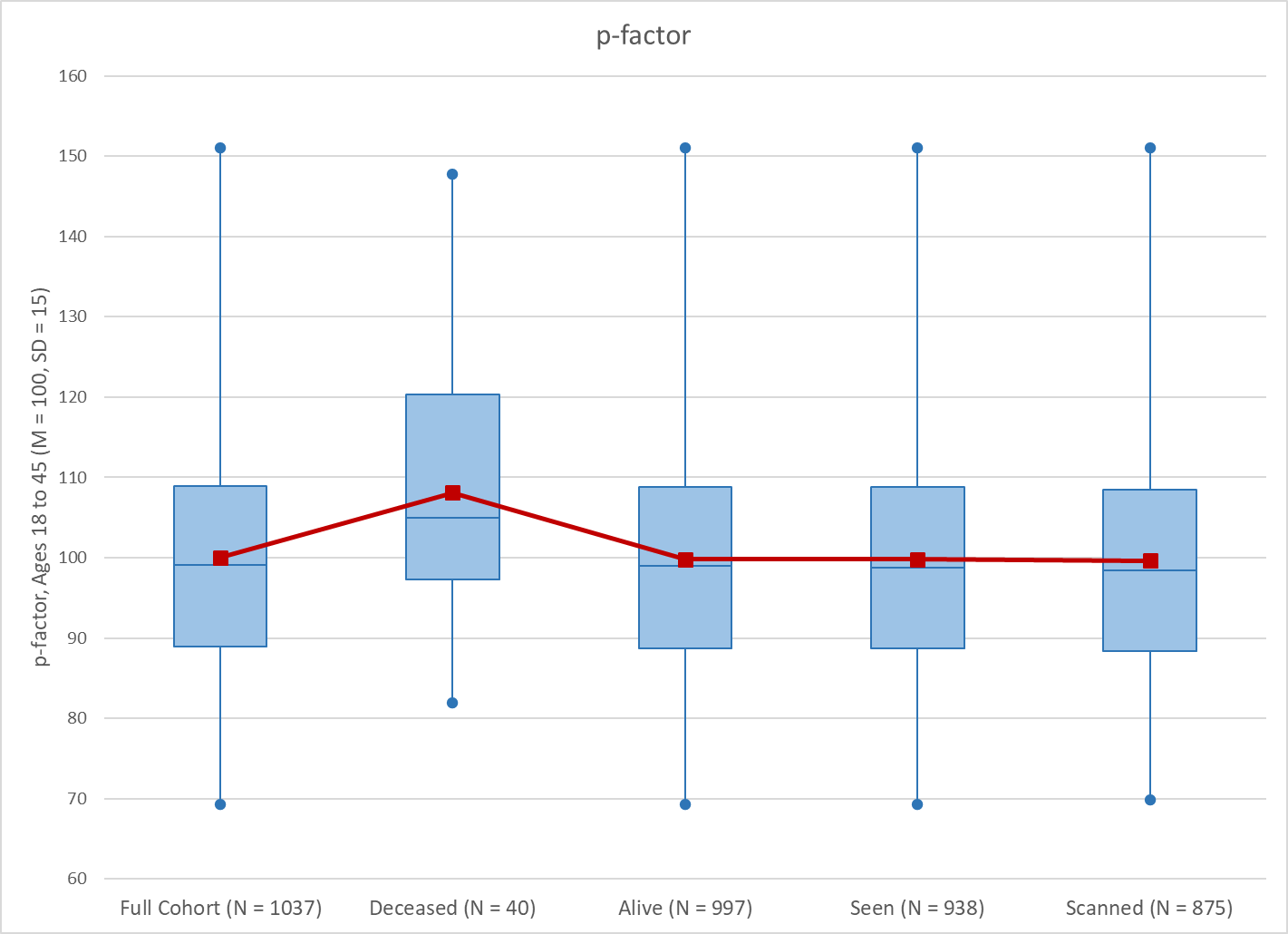

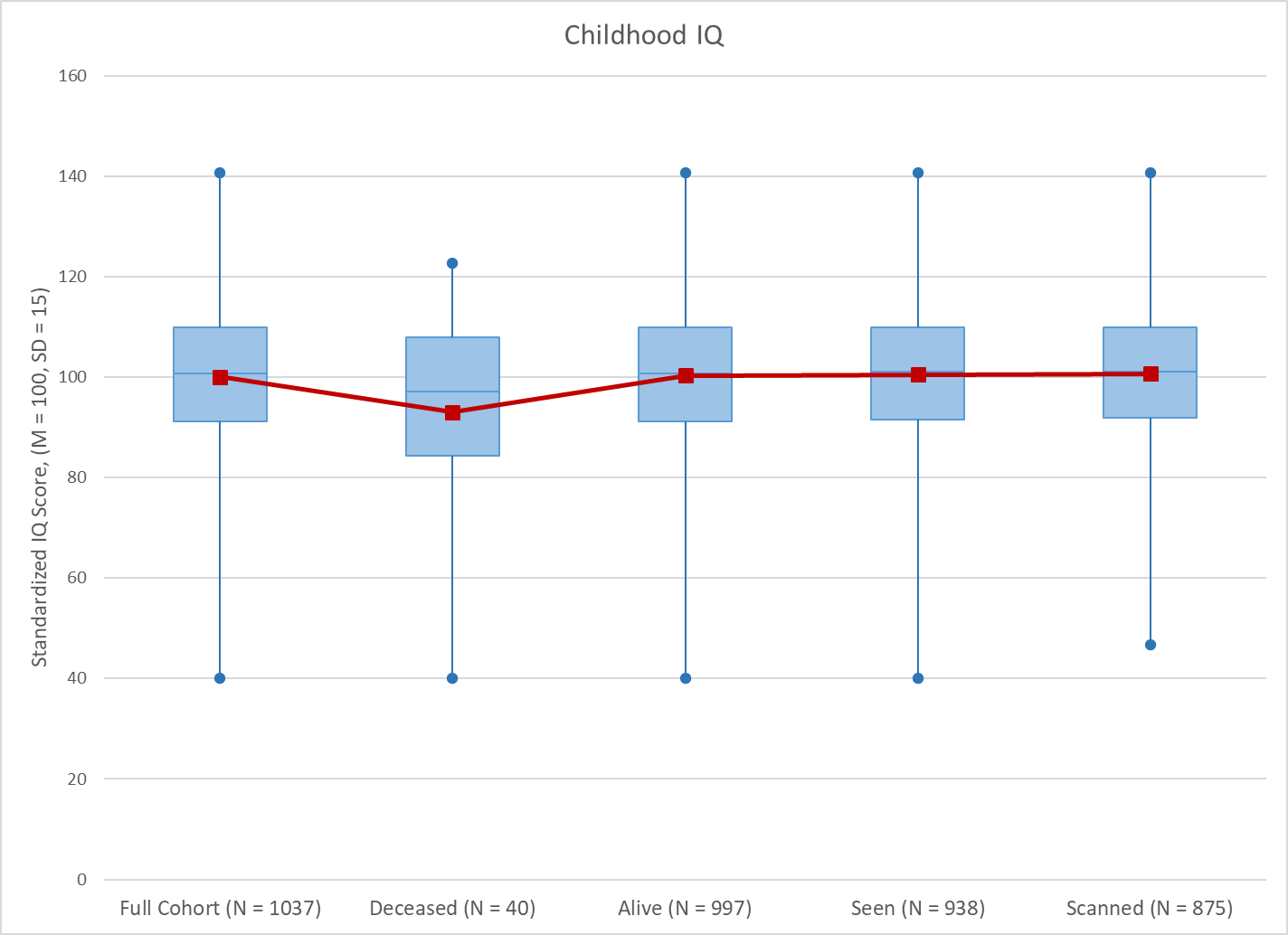

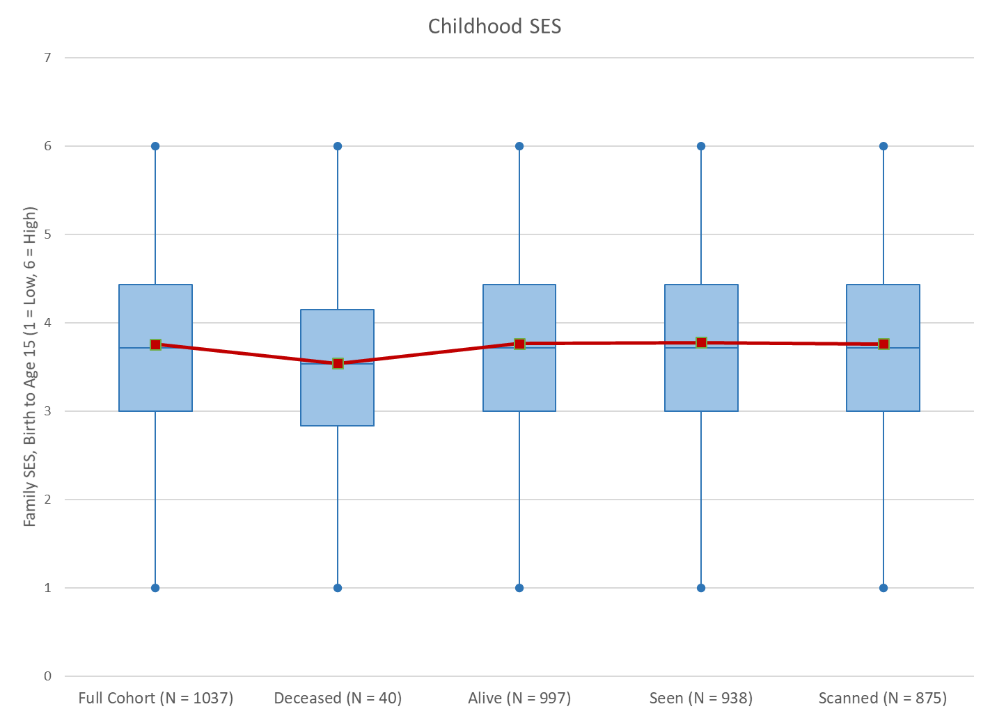

**Supplementary Figure 2.**

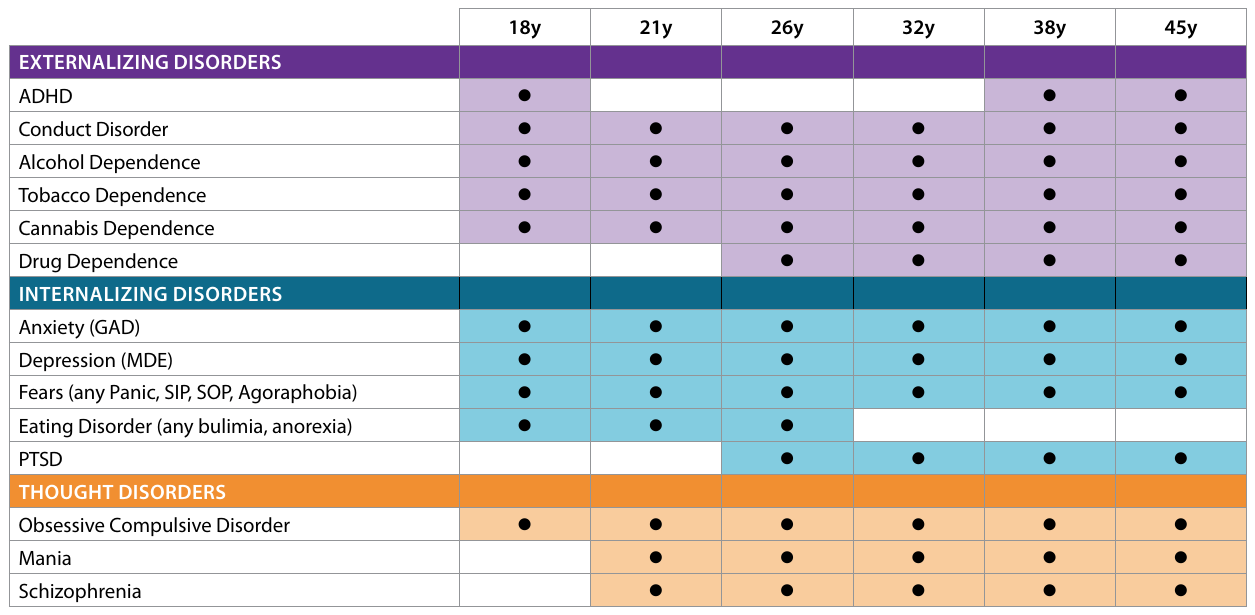

**Supplementary Figure 3**.

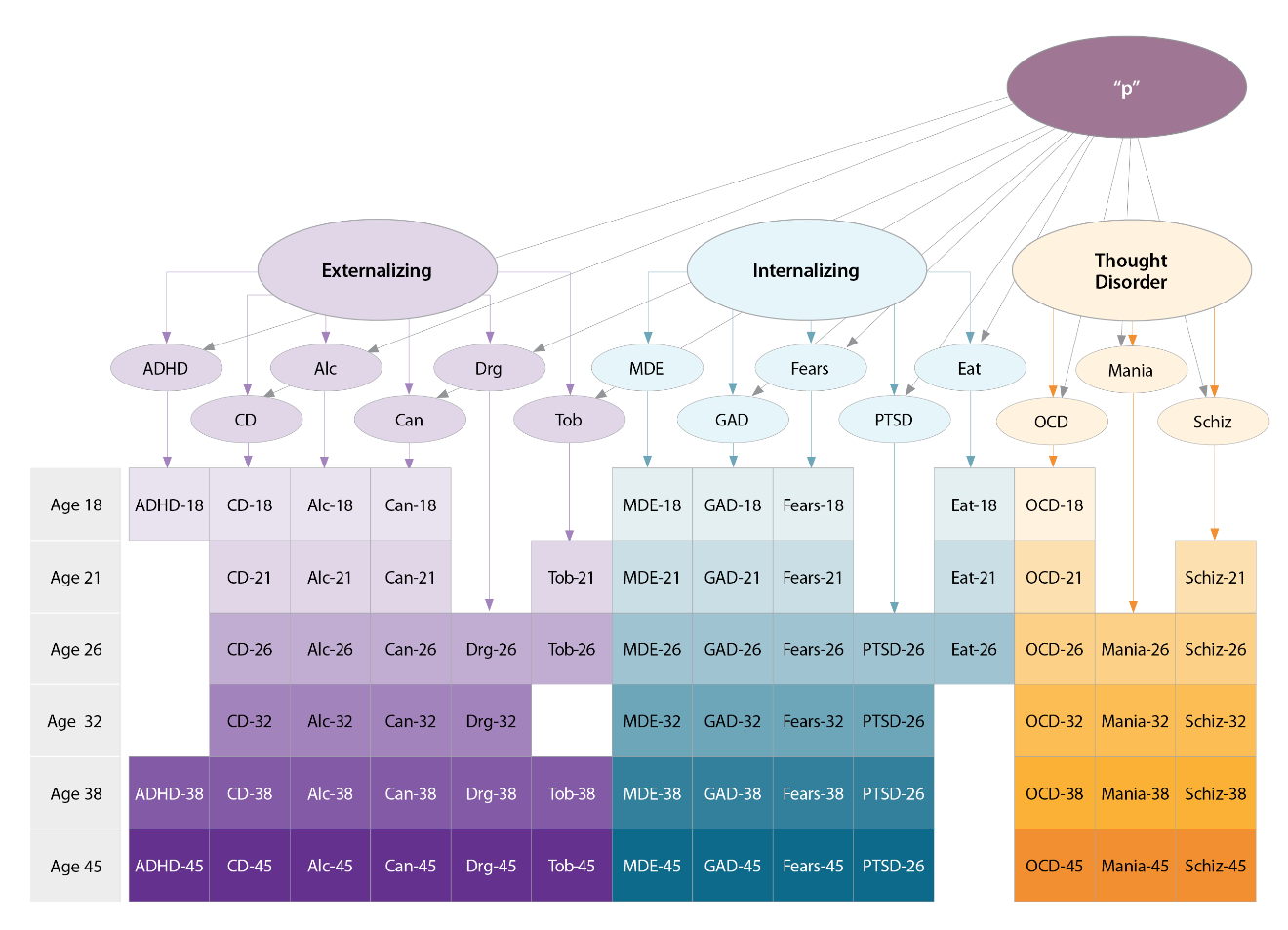

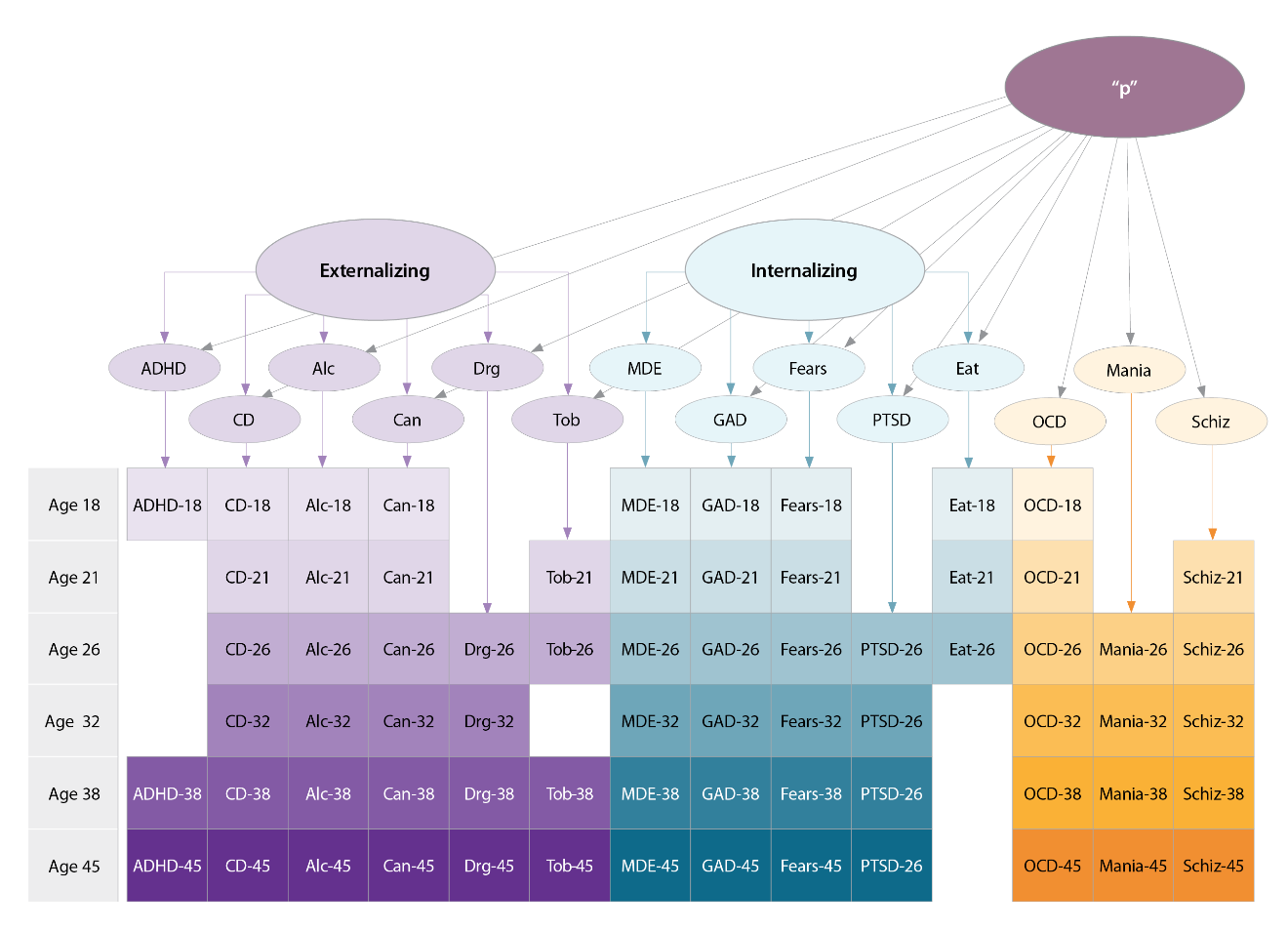

**Supplementary Figure 4.**

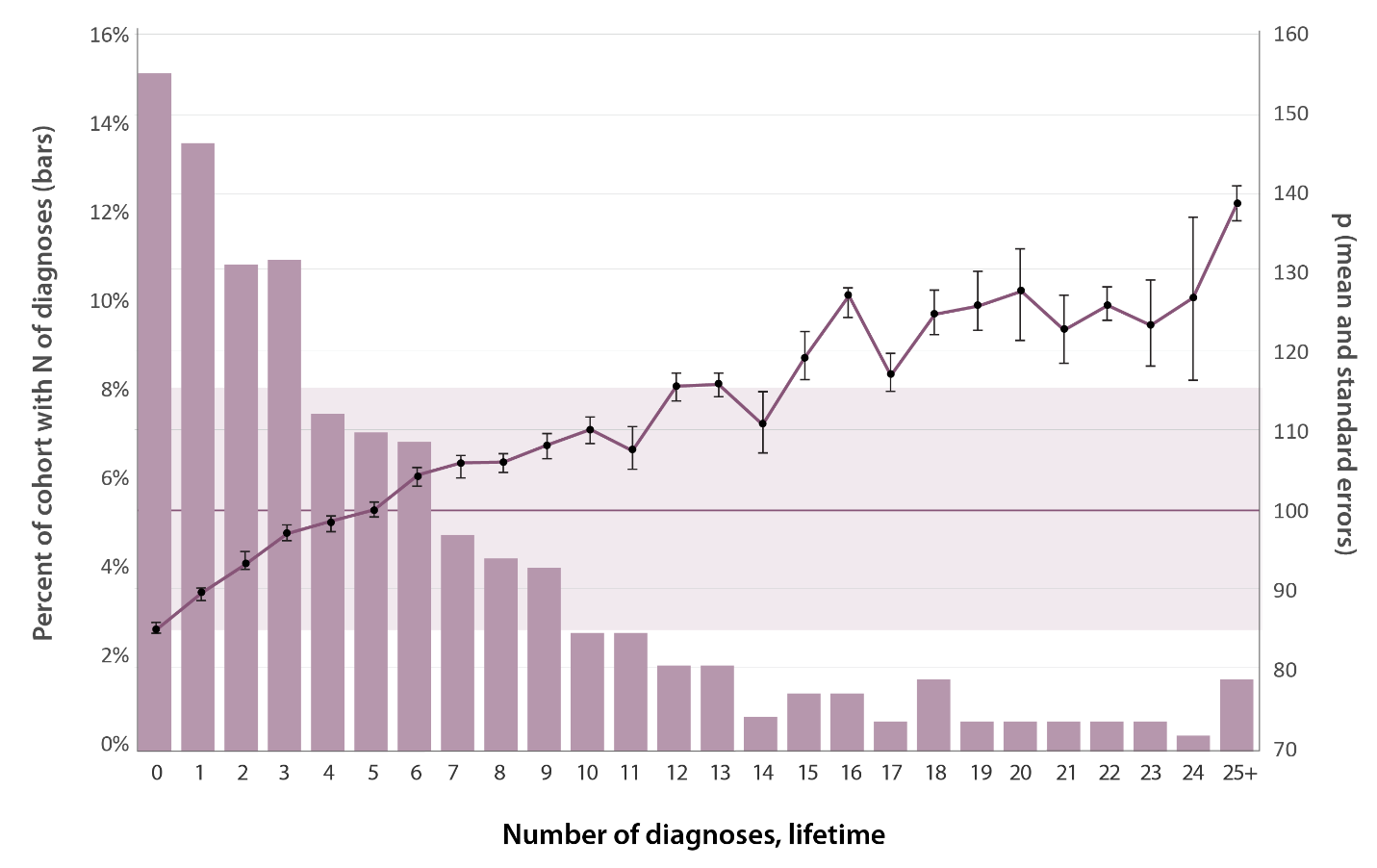

**Supplementary Table 1.** Fit statistics and standardized factor loadings for the Bifactor model.

| Model Fit Statistics | |  |  |  |  |
| --- | --- | --- | --- | --- | --- |
|  | Chi-Square (WLSMV) |  | 3695.364 | | |
|  | Degrees of Freedom |  | 2457 | | |
|  | Comparative Fit Index |  | 0.949 | | |
|  | Tucker-Lewis Index |  | 0.945 | | |
|  | RMSEA [90% CI] |  | 0.022 [0.021, 0.024] | | |
| Standardized Factor Loadings | | | p-factor | Externalizing | Internalizing |
|  | ADHD |  | 0.595 | 0.121 |  |
|  | Alcohol |  | 0.300 | 0.622 |  |
|  | Cannabis |  | 0.369 | 0.850 |  |
|  | Hard drugs |  | 0.466 | 0.694 |  |
|  | Tobacco |  | 0.450 | 0.468 |  |
|  | Conduct disorder |  | 0.504 | 0.714 |  |
|  | Major depression |  | 0.768 |  | 0.587 |
|  | Generalized anxiety |  | 0.686 |  | 0.642 |
|  | Fears/phobias |  | 0.582 |  | 0.424 |
|  | Eating disorder |  | 0.377 |  | 0.374 |
|  | PTSD |  | 0.860 |  | 0.351 |
|  | OCD |  | 0.772 |  |  |
|  | Mania |  | 0.976 |  |  |
|  | Schizophrenia |  | 0.865 |  |  |

**References**

1.	Robins LN, Cottler L, Bucholz K, Compton W. Diagnostic Interview Schedule for DSM-IV. St. Louis, MO: Washington University Press; 1995.

2.	Robins LN, Helzer JE, Croughan J, Ratcliff KS. National Institute of Mental Health Diagnostic Interview Schedule: Its history, characteristics, and validity. Arch Gen Psychiatry. 1981 Apr 1;38(4):381–9.

3.	Meier MH, Caspi A, Reichenberg A, Keefe RSE, Fisher HL, Harrington H, et al. Neuropsychological decline in schizophrenia from the premorbid to the postonset period: Evidence from a population-representative longitudinal study. Am J Psychiatry. 2014 Jan 1;171(1):91–101.

4.	APA. American Psychological Association: Diagnostic and Statistical Manual of Mental Disorders (Revised Third ed.). Washington, DC: American Psychiatric Association; 1987.

5.	APA. American Psychological Association: Diagnostic and Statistical Manual of Mental Disorders (Fourth ed.). Washington, DC: American Psychiatric Association; 1994.

6.	APA. American Psychological Association: Diagnostic and Statistical Manual of Mental Disorders (Fifth ed.). Washington, DC: American Psychiatric Association; 2013.

7.	Caspi A, Houts RM, Belsky DW, Goldman-Mellor SJ, Harrington H, Israel S, et al. The p factor: One general psychopathology factor in the structure of psychiatric disorders? Clin Psychol Sci J Assoc Psychol Sci. 2014 Mar;2(2):119–37.

8.	Heatherton TF, Kozlowski LT, Frecker RC, Fagerström KO. The Fagerström Test for Nicotine Dependence: a revision of the Fagerström Tolerance Questionnaire. Br J Addict. 1991 Sep;86(9):1119–27.

9.	Caspi A, Moffitt TE. All for one and one for all: mental disorders in one dimension. Am J Psychiatry. 2018 Apr 6;175(9):831–44.

10.	Muthen LK, Muthen BO. Mplus User’s Guide. (Eight ed.). Los Angeles, CA: Muthen & Muthen; 1998.

11.	Asparouhov T, Muthen B. Weighted Least Squares Estimation with Missing Data. Available from: Retrieved from http://www.statmodel.com/download/GstrucMissingRevision.pdf

12.	Bollen KA, Curran PJ. Latent Curve Models A Structural Equation Perspective Introduction. [Internet]. 2006. 15 p. Available from: Retrieved from <Go to ISI>://WOS:000297795700002.
